# Simple, low-cost production of DNA MS2 virus-like particles as molecular diagnostic controls

**DOI:** 10.1101/2022.09.04.506540

**Authors:** Michael A. Crone, Paul S. Freemont

## Abstract

Suitable controls are integral for the validation and continued quality assurance of diagnostic workflows. Plasmids, DNA or *in vitro* transcribed RNA are often used to validate novel diagnostic workflows, however, they are poorly representative of clinical samples. RNA phage virus-like particles packaged with exogenous RNA have been used in clinical diagnostics as workflow controls, serving as surrogates for infectious viral particles. Comparable controls for DNA viruses are more challenging to produce, with analogous DNA phages being infectious and packaging of DNA within RNA phages requiring complex purification procedures and expensive chemical linkers. We present a simple and inexpensive method to produce MS2 virus-like particles, packaged with DNA, that makes use of affinity chromatography for purification and enzymatic production of exogenous DNA suitable for packaging. The produced virus-like particles were packaged with Hepatitis B Virus DNA and were then quantified using droplet digital PCR and calibrated against the WHO international standard using a commercial assay in an accredited clinical laboratory.

---

The development of novel nucleic acid diagnostic assays for viral diseases relies on the availability of suitable materials for validation and continued quality assurance. The assessment of specificity, sensitivity, accuracy, precision and, particularly, limit of detection, requires materials that are accurately quantified and representative of the infectious agent. When available, World Health Organisation International Standards serve as the primary reference material^1^ and are often produced using clinical samples or cell cultured-derived viral particles. However, due to delays in availability, safety concerns^2^ or limited access, there is often a lag where a diagnostic assay is urgently needed and no traceable metrological standards are available. To fill this gap, interim working standards are often produced by laboratories or companies to enable proficiency testing and expedite assay development^3^ and are later calibrated against the international standard.

Working standards can be derived from a variety of different materials. For RNA viruses, *in vitro* transcribed RNA^2,4^, heat-inactivated virus particles (Zeptometrix and Qnostics) or other viral particles packaged with exogenous RNA^5,6^ have all been utilised as working standards. For DNA viruses, plasmids^7^, PCR products^8^ and viral particles packaged with exogenous DNA^9–11^ have been utilised. Although unprotected nucleic acids are popular as a part of the initial validation of diagnostic assays, alternative controls are often recommended when performing assay validation and proficiency testing within clinical laboratories^12,13^. This is because unprotected nucleic acids are degraded by nucleases when spiked into sample matrices (such as plasma or sputum) and do not serve as processing controls as they do not require extraction^9^.

Within diagnostic labs, the use of phages as validation and proficiency controls has become increasingly popular^12^. The MS2 bacteriophage, a non-enveloped virus, is widely accepted as an RNA virus surrogate and has been described for veterinary applications^14^ and as a surrogate for Influenza A^15^, SARS-CoV^15^, HIV^16^ and SARS-CoV-2^17,18^. Optimised affinity chromatography protocols have also been described, simplifying production and enabling preparation without expensive equipment^19^. Analogous DNA phages have been suggested for the preparation of “armored DNA”, but these phages can infect other bacteria^9^, suffer from low packaging efficiency^20^, are complicated to produce^11^ or are less stable in sample matrices than their RNA counterparts^13^.

In an attempt to create more stable surrogates for DNA viral particles, the MS2 RNA phage system has been successfully adapted for the packaging of exogenous DNA^21^, but the process is expensive and complex, making it unlikely to be widely adopted. Here, we describe a simplified protocol to produce MS2 virus-like particles, making use of affinity chromatography and enzymatic digestion of exogenous DNA, to enable the inexpensive manufacturing of DNA encapsulated particles. We characterise the produced particles using dynamic light scattering, qPCR, droplet digital PCR (ddPCR) and, finally, validate them using a commercial Hepatitis B virus assay within an accredited clinical laboratory.

## Results and Discussion

The MS2 bacteriophage does not natively package DNA and therefore an adapted protocol is required to package exogenous DNA sequences. Nucleic acids require a structured, sequence specific segment known as a *pac* site^5,21–23^ to be spontaneously packaged within disassembled MS2 phages. Zhang et al. have previously shown that this single stranded portion, called a translational operator DNA (TR-DNA), can be conjugated to a PCR product using chemically modified primers^22^, but their approach is complicated and expensive (Supplementary Table 3). We attempted to find a suitable enzymatic replacement to achieve the same goal, exogenous DNA that encompasses a double stranded DNA product and a single stranded 5’ *pac* site.

It has been previously reported that there are multiple enzymatic methods for the production of single stranded DNA^24^. We explored the use of λ exonuclease and T7 exonuclease to generate single-stranded DNA with the goal that two single-stranded DNA strands (complementary to one another, other than a short 5’ extension containing the *pac* site) could then be annealed successfully to generate the required DNA product. The two enzymes contrast in their digestion mechanisms, with λ exonuclease preferentially digesting phosphorylated strands, a form of negative selection, and T7 exonuclease preferentially digesting unprotected strands (strands that have not been phosphorothioated), a form of positive selection (Figure 1).

**Figure 1.**
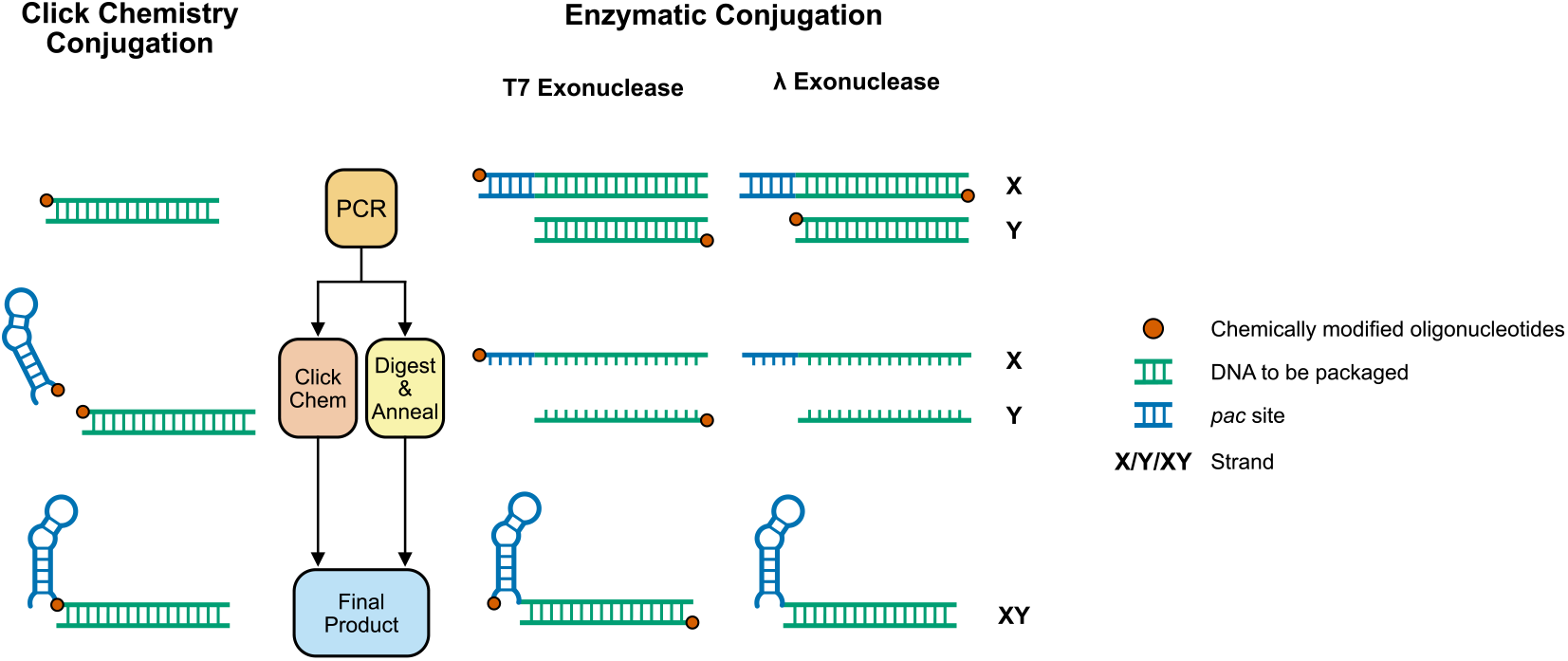
Comparison of the Click Chemistry technique used in Zhang et al.^22^ with the T7 Exonuclease and λ Exonuclease enzymatic methods. X and Y are used to denote the strand with the 5’ TR-DNA extension (X) and the shorter strand without it (Y). For the Click Chemistry approach Sulfhydryl-modified primers are used to perform the initial PCR and these are conjugated to an oligonucleotide containing the *pac* site that has an amine modification^22^. For T7 exonuclease digestion, primers are chemically modified by the addition of phosphorothioate bonds to the first 5 bases of the protected oligonucleotide. The addition of these bonds protects the strands from digestion and the final product therefore incorporates the protected oligonucleotides (positive selection). In contrast, for λ Exonuclease digestion the primers are chemically modified by phosphorylation. λ Exonuclease then preferentially digests the strands that have been phosphorylated, leaving a final product that does not incorporate the phosphorylated oligonucleotides (negative selection).

We benchmarked λ exonuclease and T7 exonuclease using two criteria: (1) the extent to which the double stranded product is digested to its single stranded form, and (2) the total yield of the final annealed product of the digested X and Y products. We showed that although it was reported that λ exonuclease did not have activity when directly added to an unpurified PCR product^24^,that the enzyme did in fact digest a double stranded PCR product to a single stranded product (Supplementary Figure 1). However, as reported previously^24^, phosphorylated oligonucleotides ordered from DNA synthesis companies are not all successfully phosphorylated, leading to only partially digested products. Since our yield of final exogenous DNA product relies on near or total digestion of the generated PCR product to its single stranded form, λ exonuclease does not perform well using the benchmark criteria.

In contrast, since T7 exonuclease digests all unprotected DNA (positive selection), there is no requirement for all oligonucleotides to have the required phosphorothioate bonds to ensure complete digestion to a single stranded product as all oligonucleotides that do not have the required bonds will be digested anyway. We show that T7 exonuclease displays activity when added directly to unpurified PCR products both with and without addition of its requisite reaction buffer (Figure 2a). We further demonstrated that X and Y PCR reactions can be combined with T7 exonuclease, digested and subsequently annealed successfully in a one pot format (workflow Figure 2b). The addition of T7 exonuclease to the X and Y PCR reactions without its accompanying reaction buffer and incubated overnight (Supplementary Figure 2 and Figure 2a) showed the best performance in terms of our benchmark criteria and were therefore the reaction conditions taken forward.

**Figure 2.**
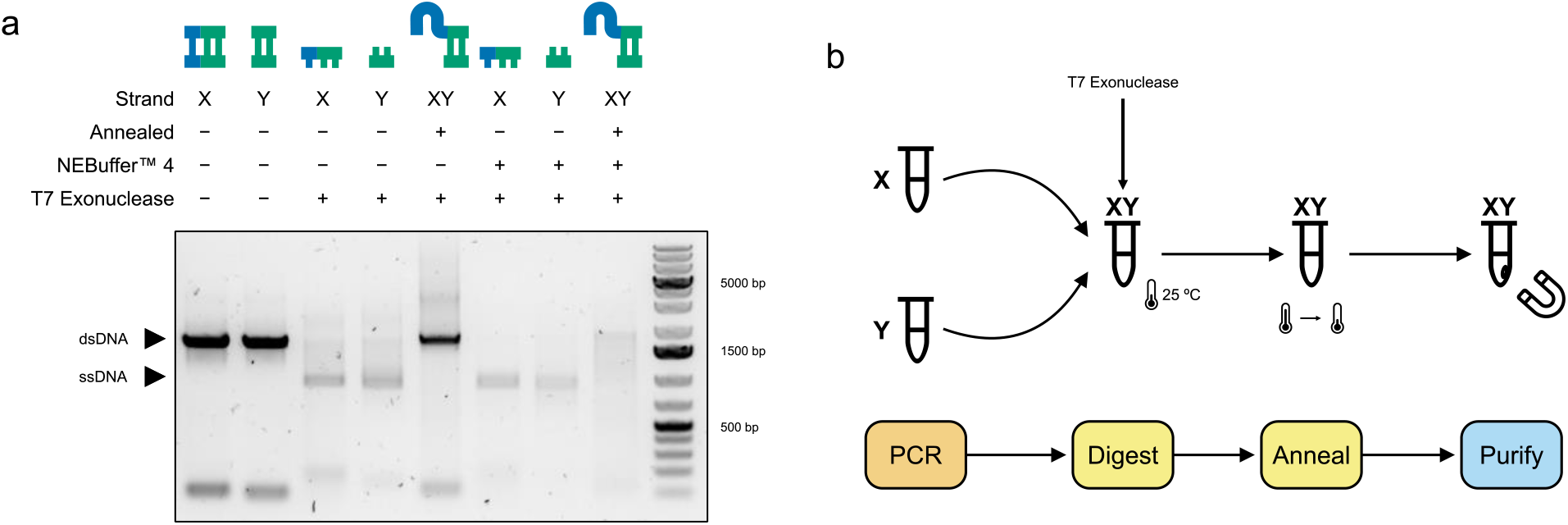
a) Agarose electrophoresis of optimisation with overnight T7 exonuclease digestion. The protected strand (X or Y or, in the case of annealed, XY) is indicated together with whether the product was annealed and the presence of reaction buffer and T7 exonuclease. T7 exonuclease showed activity both with and without addition of its reaction buffer. X and Y strands could also be combined, digested, and subsequently annealed (XY) in a single pot format. b) The final, optimised procedure for producing exogenous DNA for packaging. PCR reactions with protected X and Y strands are performed separately and then combined and supplemented with T7 exonuclease which digests the unprotected strands overnight before a final annealing step in a thermocycler. The annealed product is then purified using magnetic beads. The strand present in each tube (X, Y or XY) is shown at each step.

Using our optimised approach (Figure 2b), we amplified a portion of the Hepatitis B virus containing the X, C and S genes (Figure 3), which are targeted by several published^25,26^ and commercial assays. We then prepared the amplified product for packaging by digesting, annealing and subsequently purifying the products using magnetic beads to remove any contaminating oligonucleotides and primer dimers (Supplementary Figure 3).

**Figure 3.**
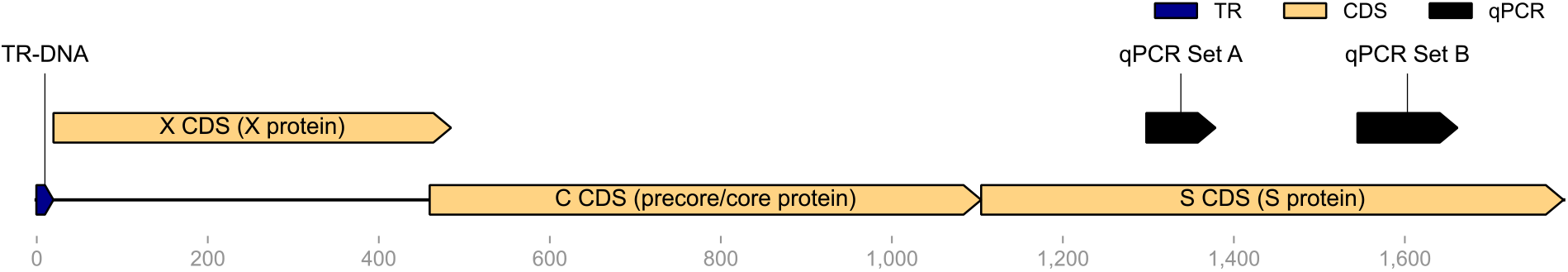
Diagram of the amplified DNA product, containing the TR-DNA and three HBV genes. The assay targets of the duplex qPCR assay (qPCR set A and B) are also displayed. Diagram was generated using DNA Features Viewer^27^.

We then proceeded to produce the viral coat protein for packaging, disassembly and subsequent reassembly (Figure 4a). We first expressed and purified unpackaged MS2 VLPs using affinity chromatography (Figure 4b) as previously described^17^. The purified particles were disassembled using glacial acetic acid as performed by Wu et al.^28^ and buffer exchanged out of glacial acetic acid and into a buffer compatible with downstream steps. The disassembled coat protein dimers were then incubated together with the exogenous DNA and subsequently purified again using affinity chromatography. Reassembly was confirmed using dynamic light scattering, showing a predominantly monodisperse peak at a diameter of approximately 30 nM (Figure 4c). Furthermore, packaging was confirmed by gel shift assay which showed a shift and more diffuse pattern of the packaged DNA when compared to the unpackaged DNA (Figure 4d)

**Figure 4.**
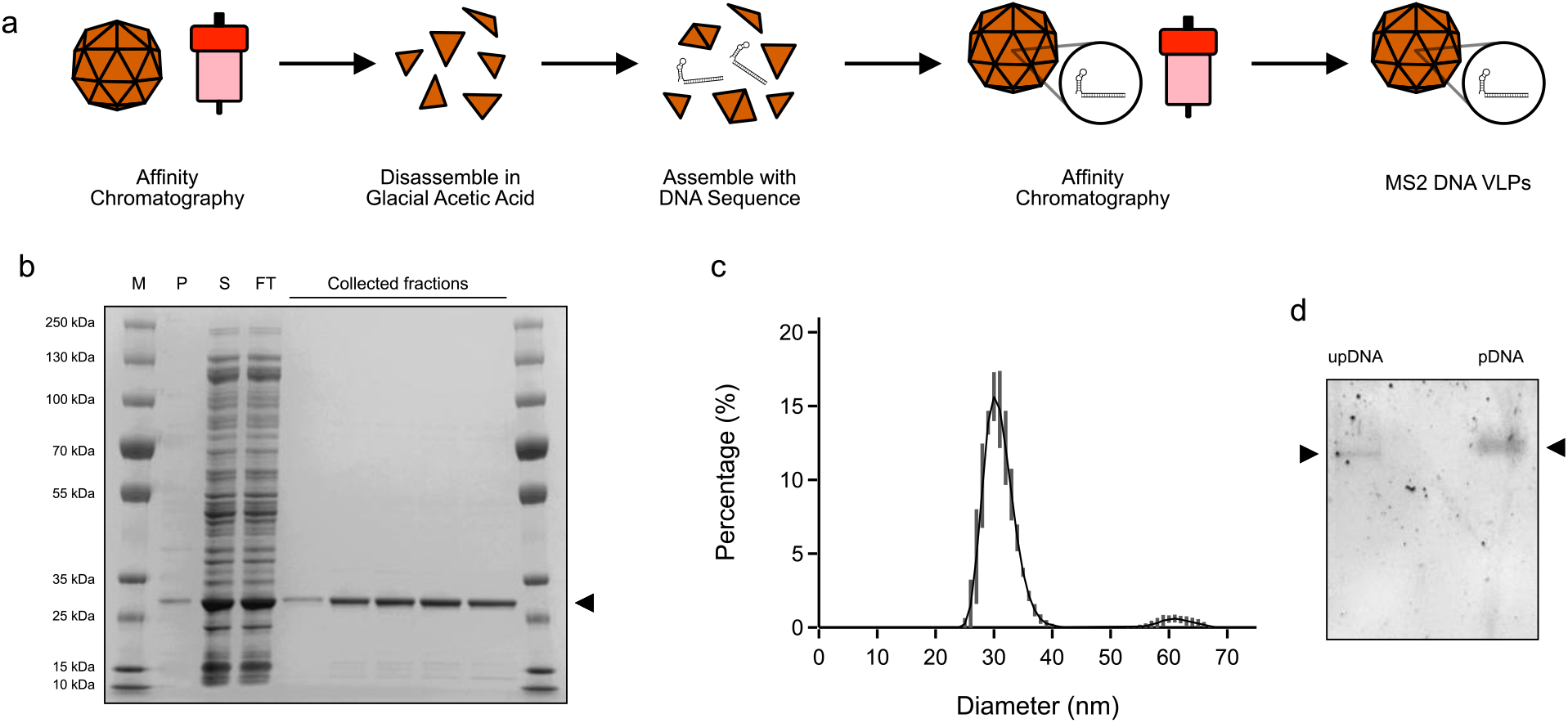
a) Preparation of DNA VLPs by protein purification of empty particles, followed by disassembly, assembly with exogenous DNA and finally a second affinity chromatography step to isolate packaged DNA VLPs. b) SDS-PAGE gel electrophoresis of the initial affinity chromatography includes a protein marker (M) followed by pellet (P) and soluble fraction (S) of the cell lysate, followed by the column flow through (FT) and collected elution fractions, where the coat protein dimer (∼ 28 kDa) is indicated by an arrow. c) Dynamic light scattering analyses of the MS2 DNA VLPs, showing a predominantly monodisperse population of ∼ 30 nM. Error bars represent the SD of three technical replicates. d) Gel shift assay showing the unpackaged DNA (upDNA) and packaged DNA within the VLP (pDNA) run side by side.

After our initial biochemical validation, we proceeded to validate the produced VLPs as molecular diagnostic controls. We first confirmed that the previously described duplex assay successfully amplified the two targets within our produced VLPs using qPCR (Supplementary Figure 4). We also used this experiment to guide the production of aliquots within a working range for further use. Then, to have an idea of the absolute concentration of the particles, we performed droplet digital PCR (ddPCR) on three individual aliquots (Figure 5). We used both primer and probe sets that we had previously validated with qPCR. Using two separate targets allowed for the determination of residual DNA found in the preparation of the VLPs. Any droplets positive for a single target were assumed to be residual DNA contamination that was not removed by DNase treatment. Concentrations are calculated using a Poisson distribution. At low concentrations the number of droplets that contain duplicate strands of DNA can be considered negligible and therefore the clustering should be representative of droplets with either residual DNA or packaged VLPs.

**Figure 5.**
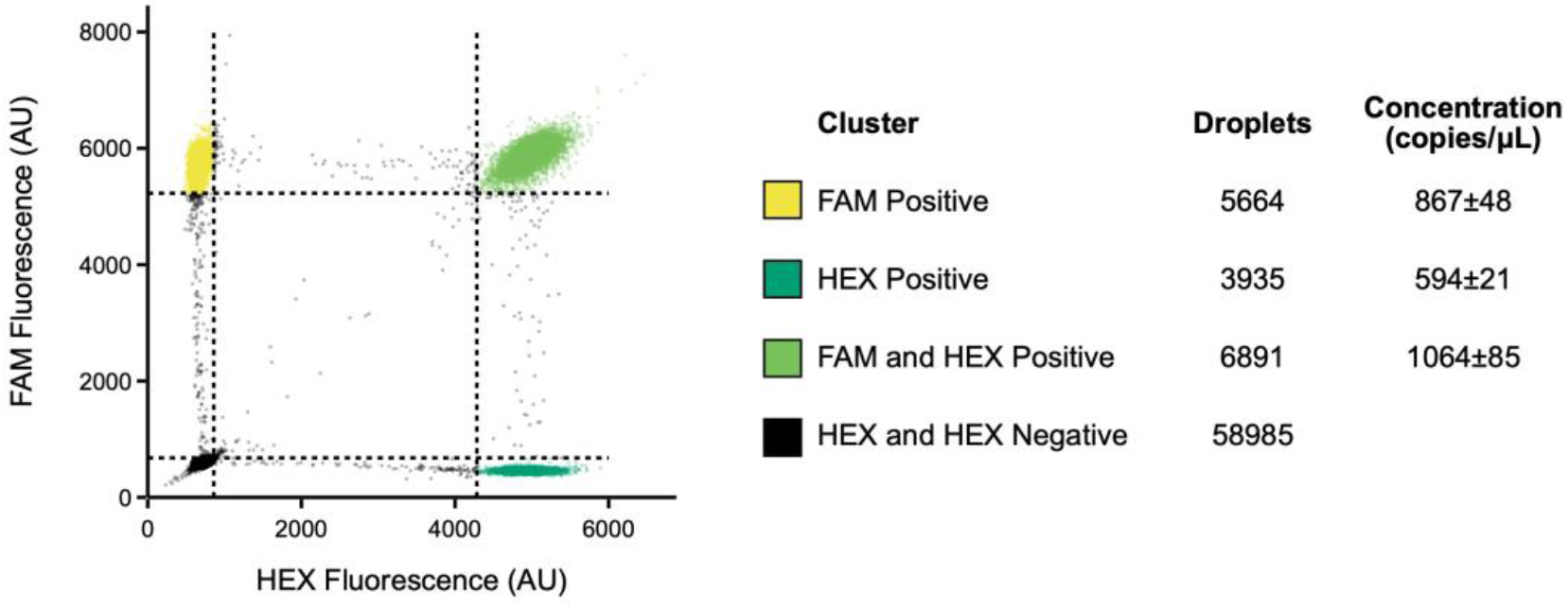
Droplet digital PCR (ddPCR) of n=3 aliquots of prepared VLPs with primer sets qPCR set A (FAM) and qPCR set B (HEX). Clustering was performed using K-Means on a single sample and the thresholds were extrapolated to the rest of the samples. Thresholds (dotted lines) are three standard deviations from the centroid of each cluster calculated using K-Means.

We were able to calculate the relative concentration of residual DNA, which was ∼ 50% for qPCR set A and ∼ 30% for qPCR set B, and the concentration of our VLPs (∼ 50% for qPCR set A and ∼ 70% for qPCR set B). The majority of the qPCR signal is therefore generated by our VLPs. Total concentration was 1931±133 copies/μL for Set A and 1658±106 copies/μL for Set B.

Finally, we validated the generated particles using a commercial assay for HBV. MS2 VLP preparations are stable when spiked into plasma^29^ and we showed consistent results over four biological replicates. Furthermore, we calibrated our developed standard against the 4^th^ WHO International Standard for HBV DNA for NAT using the commercial assay. Our working standard had a calibrated concentration of 832 IU/mL (95% PI 757-914) (Figure 6).

**Figure 6.**
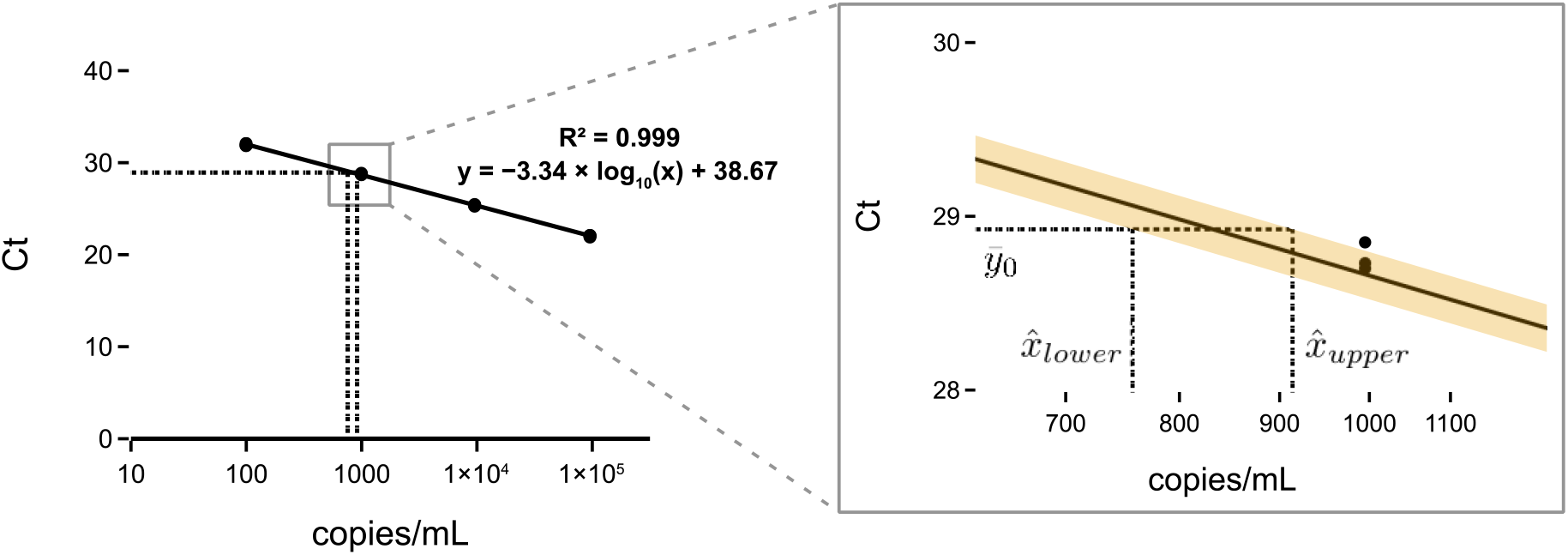
Calibration of produced VLP standard against the HBV International Standard using a Roche cobas® 6800^25^ system and the Roche cobas® HBV test. The standard curve was generated by serially diluting the 4^th^ WHO International Standard for HBV DNA for NAT. The standard curve was fit using linear regression (n=3 technical replicates of each dilution). The PCR efficiency was 99.4% (Supplementary Methods). VLP aliquots were prepared by diluting our produced VLP in human plasma. The concentration of the VLP aliquots were then calculated using the fitted equation. Error is displayed as the 95% prediction interval (PI) in yellow of n=4 biological replicates using an adaptation of Fieller’s Theorem^30^. The inset shows the mean Ct value 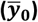 and the upper and lower bounds of the calculated VLP concentration 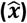.

Here we describe the development of a novel approach for the generation of dsDNA with a single stranded TR-DNA for packaging within MS2 virus-like particles. We combine this novel approach with affinity chromatography to demonstrate a simple, low-cost method for the production of DNA MS2 virus-like particles. We then validate the use of these particles as viral diagnostic controls, using a previously described duplex assay, as well as a commercial assay within a clinical laboratory. As a part of this validation we calibrate our developed standard against the 4^th^ WHO International Standard for HBV for NAT.

Suitable controls for DNA viruses are not readily available, with many companies focused on providing either diluted infectious particles or, alternatively, unpackaged DNA in the form of plasmids or chemically synthesised DNA. Furthermore, these controls are expensive and international distribution is made complicated by shipping requirements for infectious samples. We have previously shown how locally manufactured, robust and reliable diagnostic standards can play a critical role in diagnostic workflow development^17,18^. Our developed SARS-CoV-2 workflow is still making a significant contribution in diagnostic labs in London, with more than 1 500 000 tests performed by August 2022. Our previously developed SARS-CoV-2 VLPs have also been included in a Coronavirus Standards Harmonization Study as a part of the Coronavirus Standards Working Group. The only standard produced at an academic institution in the Study, it performed robustly against its commercial counterparts.

Adequate pandemic preparedness and resilience within diagnostic laboratories requires easy access to materials, not only to large commercial enterprises producing their own diagnostics for EUA or CE marking, but also to diagnostic laboratories that are developing and maintaining their own in-house assays, also known as laboratory-developed tests (LDTs), especially those in Low- and Middle-income Countries (LMICs). The reliance on commercial standards alone creates inequity in the development of clinical diagnostics. Empowering regional centres to manufacture their own standards would provide a means to democratise diagnostic assay development. By leveraging the openMTA^34^, academic institutions can provide the necessary materials and expertise, facilitating technology transfer and allowing for the decentralised production of biosafe, “open source” standards, such as virus-like particles, that rely only on the availability of low-cost protein purification equipment^17^.

We believe that by providing a toolkit for the development of reliable, robust and quantitative standards for RNA and DNA viruses and making it available under the openMTA we can begin to enable laboratories around the world to develop their own diagnostic assays. Here we add the DNA MS2 VLP to this toolkit to complement our previously developed method for making low-cost RNA MS2 VLPs. We have made all required materials available under the openMTA to enable both commercial and academic institutions to work with the materials without limitations.

## Methods

### Primers and probes and plasmid to produce viral DNA sequence

Primers and probes were ordered from IDT and can be found in Supplementary Table 1 and 2. A construct containing the X, C and S genes of the Hepatitis B virus (accession number: KY003230^35^) was ordered from GeneArt (ThermoFisher Scientific) and is available from Addgene (Addgene # 179155).

### Production of partially single stranded DNA for packaging using T7 Exonuclease

Two sets of PCR reactions were setup to create the exogenous DNA for packaging. The X PCR was performed using a phosphorothiated forward primer and unprotected reverse primer and the Y PCR was performed with an unprotected forward primer and phosphorothiated reverse primer. PCR reactions were performed with a final primer concentration of 1 mM using Q5® High-Fidelity 2X Master Mix (NEB). Equal volumes of X and Y PCR reactions were then combined, supplemented with 0.2 U/μL of T7 Exonuclease (NEB) and incubated at 25 °C overnight before being heated to 95 °C and slowly annealed (−0.1 °C/sec) in a thermocycler.

Digested, annealed exogenous DNA was concentrated in an Amicon® Ultra 0.5mL 30K Centrifugal Filter (Merck Millipore) and two washes were performed with Invitrogen TE Buffer (ThermoFisher Scientific). The final concentrated product was made up to 100 μL with Invitrogen TE Buffer (ThermoFisher Scientific) and 0.7x AMPure XP beads (Beckman Coulter) were added and the product purified according to the manufacturer’s instructions (two washes with 80% ethanol were performed and the final product was eluted in UltraPure™ DNase/RNase-Free Distilled Water (ThermoFisher Scientific)). Eluted DNA was diluted and quantified using the Qubit™ dsDNA HS kit (ThermoFisher Scientific) on a Qubit™ 3 Fluorometer (ThermoFisher Scientific).

### Expression, purification and disassembly of MS2 coat protein dimers and maturation protein

A plasmid construct containing only the maturation protein and his-tagged coat protein dimer (Addgene # 179156) was transformed into Rosetta2™ (DE3) pLysS cells (Merck). An overnight culture was used to inoculated 200 mL of Terrific Broth (Merck) supplemented with 50 mg/mL of Kanamycin (Merck), and grown at 30 °C, 200 r.p.m. until an OD of 0.8. The culture was induced by supplementing with 0.5 mM IPTG (Merck) and grown at 30 °C for a further 16 h. Cells were collected at 3220 × g at 4 °C and stored at − 20 °C for later purification.

All protein purification steps were performed at 4 °C. The cell pellet was resuspended in 4 mL Sonication Buffer (50 mM Tris-HCl pH 8.0, 5 mM MgCl_2_, 5 mM CaCl_2_, and 100 mM NaCl) with 700 U RNase A (Qiagen), 2500 U BaseMuncher Endonuclease (Abcam), and 200 U TURBO DNase (ThermoFisher Scientific). The cells were sonicated for a total of 2 min (50% amplitude, 30 s on, 30 s off) on wet ice. The lysate was then incubated for 3 h at 37 °C. The lysate was centrifuged at 10,000 × g for 10 min at room temperature in a microcentrifuge. The supernatant was then filtered with a Minisart® 5 μ m cellulose acetate (SFCA) filter (Sartorius) before being mixed 1 : 1 with 2× Binding Buffer (100 mM monosodium phosphate monohydrate pH 8.0, 30 mM Imidazole, 600 mM NaCl).

Supernatant was applied to a 5 mL HiTrap® TALON® Crude column (Cytiva) on an ÄKTA pure (Cytiva) primed with Binding Buffer (50 mM monosodium phosphate monohydrate pH 8.0, 15 mM Imidazole, 300 mM NaCl). The protein was eluted with elution buffer (50 mM monosodium phosphate monohydrate pH 8.0, 200 mM Imidazole, 300 mM NaCl) and then desalted and buffer exchanged into STE buffer (10 mM Tris-HCl pH 7.5, 1 mM EDTA, 100 mM NaCl) using an Amicon® Ultra-15 10 K Centrifuge Filter (Merck Millipore). The protein concentration was measured using the Qubit Protein Assay Kit and Qubit 3 Fluorometer (Thermo Fisher Scientific).

The purified protein was incubated with cold glacial acetic acid (final concentration 66% V/V) for 30 minutes on ice before being centrifuged at 6600 × g for 20 minutes at 4 °C. The protein sample was then buffer exchanged into Sonication Buffer (50 mM Tris-HCl pH 8.0, 5 mM MgCl2, 5 mM CaCl2, and 100 mM NaCl) and concentrated down to 500 mL using an Amicon® Ultra-15 10 K Centrifuge Filter (Merck Millipore).

### Encapsulation of DNA within MS2 virus-like particles and final purification

Purified exogenous DNA was incubated in a Protein LoBind tube (Eppendorf) with a 10-fold molar excess of disassembled MS2 coat protein dimers for 3 hours at room temperature before being incubated for 36 hours at 4 °C. Sonication buffer was added up to 1.5 mL, supplemented with 100 U of TURBO DNase (ThermoFisher Scientific) and incubated for 90 minutes at 37 °C.

The sample was then mixed 1 : 1 with 2× Binding Buffer (100 mM monosodium phosphate monohydrate pH 8.0, 30 mM Imidazole, 600 mM NaCl) and was applied to a 5 mL HiTrap® TALON® Crude column (Cytiva) on an ÄKTA pure (Cytiva) primed with Binding Buffer (50 mM monosodium phosphate monohydrate pH 8.0, 15 mM Imidazole, 300 mM NaCl). The protein was eluted with elution buffer (50 mM monosodium phosphate monohydrate pH 8.0, 200 mM Imidazole, 300 mM NaCl) and then desalted and buffer exchanged into STE buffer (10 mM Tris-HCl pH 7.5, 1 mM EDTA, 100 mM NaCl) using an Amicon® Ultra-15 30 K Centrifuge Filter (Merck Millipore).

### Dynamic Light Scattering

DLS was performed using a Zetasizer Nano (Malvern Panalytical) according to the manufacturer’s instructions.

### Gel Shift Assay

The gel shift assay was run as a 1% agarose gel and post stained with SYBR Gold nucleic acid stain (ThermoFisher Scientific).

### qPCR

Quantitative PCR was performed using a previously described duplex assay^25^ with the primers and probes found in Supplementary Table 2. VLPs were lysed by heating to 95 °C for 5 minutes in a thermocycler before being added to qPCR reactions. qPCR reactions (20 μL) used TaqPath™ qPCR Master Mix, CG (ThermoFisher Scientific) with final primer and probe concentrations of 400 nM and 200 nM and a sample volume of 1 μL. The reaction was then thermocycled (50 °C for 2:00, 95 °C for 5:00 and 45 cycles of 95 °C for 0:10 and 60 °C for 1:00) on a BioRad CFX96 qPCR machine.

### ddPCR

Droplet Digital PCR (ddPCR) was performed using a previously described duplex assay^25^ with the primers and probes found in Supplementary Table 2. VLPs were lysed by heating to 95 °C for 5 minutes in a thermocycler before being added to ddPCR reactions. ddPCR reactions (20 μL) used ddPCR™ Multiplex Supermix (Bio Rad) with final primer and probe concentrations of 900 nM and 250 nM and a sample volume of 2 μL. Droplets were then generated using a QX200™ Droplet Generator (Bio Rad) and transferred to a PCR plate and sealed, all according to the manufacturer’s instructions. The reaction was then thermocycled (95 °C for 10:00, 40 cycles of 94 °C for 0:30 and 60 °C for 1:00 and a final enzyme inactivation step of 98 °C for 10:00) on a BioRad C1000 touch PCR machine. Finally, droplets were read using the QX200™ Droplet Reader (Bio Rad). Data was then analysed using a python implementation (https://github.com/mcrone/plotlydefinerain) of an online tool called Defining The Rain (https://definetherain.org.uk/) with added support for two colour channels.

### Calibration against WHO International Standard using a Commercial Assay

The 4th WHO International Standard for HBV DNA for NAT was ordered from NIBSC. The Standard was resuspended according to the instructions for use. Calibration curve samples were generated by diluting the International Standard with human plasma (Merck) to create a standard curve ranging from 955 000 IU/mL to 95.5 IU/mL with three technical replicates of each concentration. Aliquots of our working standard were generated by spiking 50 μL of our generated VLPs into 950 μL of human plasma (Merck). The calibration curve samples and the aliquots of working standard were then tested using a cobas® 6800 system (Roche) and cobas® HBV Test (Roche) according to the manufacturer’s instructions. The working standard calibration was performed by fitting of the standard curve using linear regression and the error was calculated using a statistical tool^30^ based on Fieller’s theorem^36^. Further details are available in the Supplementary Methods. Residuals of the linear regression can be visualised in Supplementary Figure 5. A python implementation of the calculation of the prediction intervals and plots is available on GitHub (https://github.com/mcrone/qpcr_abs_calibration).

## Acknowledgements

This work is supported by the UK Dementia Research Institute which receives its funding from UK DRI Ltd, funded by the UK Medical Research Council, Alzheimer’s Society and Alzheimer’s Research UK. We also acknowledge funding from UKRI-EPSRC (EP/R014000/1) and Community Jameel. We thank Professor Charles Bangham for the use of his digital PCR machine. We thank Dr Aileen Rowan for supervising the preparation of the international standard samples under BSL3 conditions. We thank Prof. Graham P Taylor, Dr Marcus Pond and Dr Paul Randell for providing feedback on the manuscript and we thank Hitesh Mistry, Pinglawathee Madona and Louise Shelley for assistance in running the Roche assay.

## Author Contributions

M.A.C. performed experiments, analysed and interpreted the data. P.F. supervised the work. P.F. and M.A.C. made substantial contributions to the conception and design of the work and contributed to the draft of the work.

