## Supplementary Materials for "Simple, low-cost production of DNA MS2 virus-like particles as molecular diagnostic controls"

#### Supplementary Tables

**Supplementary Table 1.** Oligonucleotides required for exogenous DNA amplification

| Process | Forward Sequence | Reverse Sequence |
|---|---|---|
| Exogenous DNA [T7 Exonuclease] | A*A*C*A*T*GAGGATTACCCATGTATGGCTGCTAGGCTGTACTGC | TTAAATGTATACCCAGAGACAAAAGAAAATTG |
| | ATGGCTGCTAGGCTGTACTGC | T*T*A*A*A*TGTATACCCAGAGACAAAAGAAAATTG |
| Exogenous DNA [Lambda Exonuclease] | AACATGAGGATTACCCATGTATGGCTGCTAGGCTGTACTGC | /5Phos/TTAAATGTATACCCAGAGACAAAAGAAAATTG |
| | /5Phos/ATGGCTGCTAGGCTGTACTGC | TTAAATGTATACCCAGAGACAAAAGAAAATTG |

**Supplementary Table 2.** Oligonucleotides required for qPCR and ddPCR quantification (derived from a previously published assay[1]).

| Set | Forward Sequence | Reverse Sequence | Probe Sequence |
|---|---|---|---|
| A | GTCCTCCAATTTGTCCTGG | TGAGGCATAGCAGCAGGAT | /56-FAM/CTGGATGTG/ZEN/TCTGCGGCGTTTTATCAT/3IABkFQ/ |
| B | CACCTGTATTCCCATCCCATC | AGCCCTACGAACCACTGAACA | /5HEX/AAACGGACT/ZEN/GAGGCCCACTCCCA/3IABkFQ/ |

**Supplementary Table 3.** Cost comparison between the method described by Zhang et al. and the method described in this study for the production of a single DNA VLP. Costs of common consumables like pipette tips and chemicals used to make buffers have been omitted. Pricing is current as of August 2022, but could be subject to change.

| | | | | | Zhang et al.[2] | This Study |
|---|---|---|---|---|---|---|
| **Stage** | **Item** | **Manufacturer** | **Catalog Number** | **List Price** | **Per VLP** | **Per VLP** |
| Exogenous DNA Production | DNA Synthesis (1000 bp) | Twist Bioscience | | £50 | £50 | £50 |
| | Oligo Synthesis | IDT | | | £100 | £45 |
| | Q5® High-Fidelity 2X Master Mix | NEB | M0492S | £134 | £21 | £43 |
| | Amicon® Ultra 0.5mL 3kDa | Merck Millipore | UFC500308 | £46.60 | £5.83 | |
| | Amicon® Ultra 0.5mL 50kDa | Merck Millipore | UFC505008 | £46.60 | £11.65 | |
| | Zeba™ Spin Desalting Columns, 40K MWCO, 0.5 mL | ThermoFisher Scientific | 87766 | £154.00 | £6.16 | |
| | T7 Exonuclease | NEB | M0263S | £57 | | £18 |
| | AMPure XP | Beckman Coulter | A63880 | £230.90 | | £3.23 |
| | Amicon® Ultra 0.5mL 30kDa | Merck Millipore | UFC503008 | £46.60 | | £5.83 |
| | TE Buffer | ThermoFisher Scientific | 12090015 | £47 | £0.47 | £0.47 |
| | | | | *Total DNA production* | £195.54 | £166 |
| Protein Purification | TURBO™ DNase | ThermoFisher Scientific | AM2238 | £125 | £18.75 | £18.75 |
| | RNase A | Qiagen | 19101 | £225 | £9.00 | £9.00 |
| | Basemuncher | Abcam | ab270049 | £155 | £15.50 | £15.50 |
| | HiPrep 16/60 Sephacryl S-200 HR | Cytiva | 17116601 | £585 | £585 | |
| | HiLoad 16/600 Superdex 75 pg | Cytiva | 28989333 | £1,904 | £1,904 | |
| | SnakeSkin™ Dialysis Tubing, 10K MWCO, 22 mm | ThermoFisher Scientific | 68100 | £184 | £5 | |
| | Minisart® Syringe Filter, SFCA, Pore Size 5 mm | Sartorius | 17594 | £848 | | £1.70 |
| | HiTrap TALON crude | Cytiva | 28953767 | £510 | | £102 |
| | Amicon® Ultra-15 10 kDa (2) | Merck Millipore | UFC901008 | £97.90 | | £24.48 |
| | Amicon® Ultra-15 30 kDa | Merck Millipore | UFC903008 | £97.90 | | £12.24 |
| | | | | *Total Protein Purification* | £2,537.36 | £183.71 |
| | | | | **Total Cost** | **£2,732.90** | **£349.36** |

**Supplementary Table 4.** Raw Roche® cobas 6800 results for the diluted International Standard and VLP samples using the Roche cobas® HBV assay.

| Sample | Target 1 Ct | Measured IU/mL | QS Ct | QS Result | Valid |
|---|---|---|---|---|---|
| 95500 | 21.92 | 107000 | 32.43 | Valid | Yes |
| 95500 | 22.08 | 98200 | 32.46 | Valid | Yes |
| 95500 | 22.09 | 85100 | 32.26 | Valid | Yes |
| 9550 | 25.28 | 15100 | 32.93 | Valid | Yes |
| 9550 | 25.45 | 8390 | 32.24 | Valid | Yes |
| 9550 | 25.4 | 9140 | 32.32 | Valid | Yes |
| 995 | 28.73 | 940 | 32.33 | Valid | Yes |
| 995 | 28.7 | 1010 | 32.41 | Valid | Yes |
| 995 | 28.85 | 1030 | 32.58 | Valid | Yes |
| 99.5 | 32.09 | 98 | 32.4 | Valid | Yes |
| 99.5 | 31.86 | 115 | 32.4 | Valid | Yes |
| 99.5 | 31.88 | 147 | 32.77 | Valid | Yes |
| VLP | 28.81 | 932 | 32.4 | Valid | Yes |
| VLP | 28.8 | 835 | 32.23 | Valid | Yes |
| VLP | 29.13 | 908 | 32.68 | Valid | Yes |
| VLP | 28.96 | 756 | 32.25 | Valid | Yes |
| Blank | - | Target Not Detected | 32.41 | Valid | Yes |

### Supplementary Figures

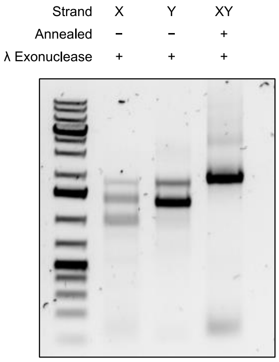

**Supplementary Figure 1. Agarose Electrophoresis showing digestion with λ Exonuclease. λ Exonuclease was added directly to the completed PCR reactions without reaction buffer.**

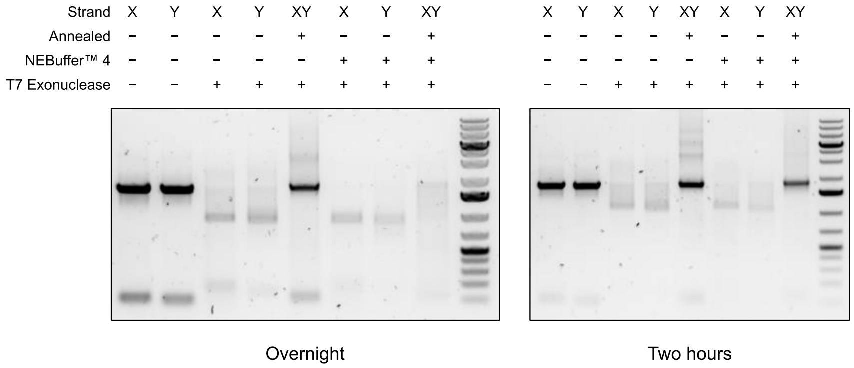

**Supplementary Figure 2. Agarose Electrophoresis showing optimisation of digestion conditions with T7 Exonuclease. Reactions were tested with and without buffer, individually and after adding F and R together and with a two hour digestion or overnight at 25 °C.**

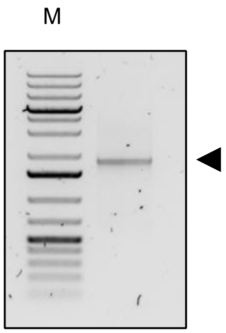

**Supplementary Figure 3. Agarose Electrophoresis with DNA Ladder (M) and magnetic bead purified product after digestion and annealing (indicated with the arrow).**

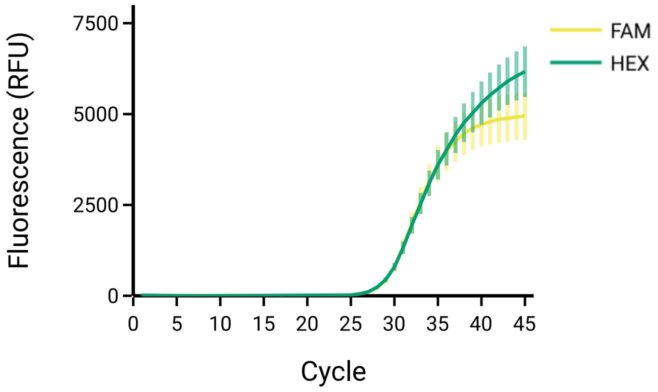

**Supplementary Figure 4. Raw qPCR curves of DNA MS2 VLPs detected with the duplex assay[1]. Error bars represent the SD of n = 5 technical replicates.**

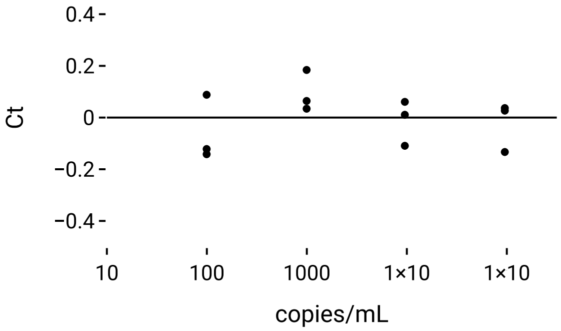

**Supplementary Figure 5. Residuals obtained after fitting the standard curve using linear regression (SEq 1).**

#### Supplementary Methods

##### *Production of partially single stranded DNA for packaging using $\lambda$ Exonuclease*

Two sets of PCR reactions were setup to create the exogenous DNA for packaging. The X PCR was performed using a phosphorylated reverse primer and unmodified forward primer and the Y PCR was performed with a phosphorylated forward primer and unmodified reverse primer. PCR reactions were performed with a final primer concentration of 1 μM using Q5® High-Fidelity 2X Master Mix (NEB). Equal volumes of X and Y PCR reactions were then combined, supplemented with 0.2 U/μL of λ Exonuclease (ThermoFisher Scientific) and incubated at 37 °C for 30 minutes before being heated to 95 °C and slowly annealed (-0.1 °C/sec) in a thermocycler.

##### *Concentration and error calculation for working standard calibration*

The working standard calibration was performed using linear regression and the error was calculated using a stastistical tool[3] based on Fieller's theorem[4]. For completeness the equations below are included as found in Pizzamiglio et al[3].

The data from the standard curve was initially fit by linear regression to SEq 1. The residuals obtained using the fitted equation can be visualized in Supplementary Figure 5.

$$y_{ij} = \beta_0 + \beta_1 x_i$$ **SEq 1**

where $y_{ij}$ specifies the value of the Ct values determined for the j-th replicate (j = 1,2,...,Ji ) at log $x_i$ different (i = 1,2,...,I) WHO international standard dilutions.

PCR efficiency could then be calculated using SEq 2:

$$\text{Efficiency} = 10^{(-1/\beta_1)} - 1$$ **SEq 2**

The estimate of the unknown log starting concentration ($\hat{x}_0$) of the unknown VLP concentration ($x_0$) is then calculated by substituting the mean Ct of the replicates ($\bar{y}$) of the VLP working standard into SEq 3.

$$\hat{x}_0 = \frac{\bar{y} - \beta_0}{\beta_1}$$ **SEq 3**

The variance for the standard was then estimated using SEq 4.

$$s_p^2 = \frac{\sum_{i=1}^{I} \sum_{j=1}^{J_i} (y_{ij} - \bar{y}_i)^2}{\sum_{i=1}^{I} (J_i - 1)}$$ **SEq 4**

where $\bar{y}_i$ is the mean of the Ct values at the i-th standard dilution

We can then define the deviance ($s_{xx}$) of the $x_i$ values as SEq 5:

$$s_{xx} = \sum_{i=1}^{I} J_i (x_i - \bar{\bar{x}})^2$$ **SEq 5**

And the mean ($\bar{\bar{x}}$) of the $x_i$ values as SEq 6:

$$\bar{\bar{x}} = \frac{\sum_{i=1}^{I} J_i x_i}{\sum_{i=1}^{I} J_i}$$ **SEq 6**

$t_{f;1-\alpha/2}$ is the value which corresponds to a critical value where the significance is α and there are $f$ degrees of freedom.

The confidence intervals of $x_0$ are obtained by calculating the roots of the quadratic equation SEq 7.

$$Ax^2 + 2Bx + C = 0$$ **SEq 7**

where:

$$A = \beta_1^2 - \frac{s_p^2}{s_{xx}} t_{f;1-\alpha/2}^2$$ **SEq 8**

$$B = \beta_0\beta_1 - \bar{y}\beta_1 + \frac{s_p^2 \bar{\bar{x}}}{s_{xx}} t_{f;1-\alpha/2}^2$$ **SEq 9**

$$C = \bar{y}^2 + \beta_0^2 - 2\bar{y}\beta_0 - \frac{s_p^2}{K} t_{f;1-\alpha/2}^2 - \left( \frac{s_p^2 \sum_{i=1}^{I} J_i x_i^2}{s_{xx} \sum_{i=1}^{I} J_i} \right) t_{f;1-\alpha/2}^2$$ **SEq 10**

finally, by defining $g$ as:

$$g = \frac{s_p^2 t_{f;1-\alpha/2}^2}{s_{xx} \beta_1^2}$$ **SEq 11**

We can obtain the two roots of SEq 7 and the required confidence limits of $x_0$ using SEq 12:

$$\left.\begin{matrix} \hat{x}_{upper} \\ \hat{x}_{lower} \end{matrix}\right\} = \hat{x}_0 + \frac{(\hat{x}_0 - \bar{\bar{x}})g \pm (s_p t_{f;1-\alpha/2}/\beta_1)\{[(\hat{x}_0 - \bar{\bar{x}})^2/s_{xx}] + (1-g)(1/K + 1/n)\}^{\frac{1}{2}}}{1-g}$$ **SEq 12**

The prediction limits of the Ct value of the VLP sample ($\bar{y}$) are then calculated using SEq 13:

$$\left.\begin{matrix} \hat{y}_{upper} \\ \hat{y}_{lower} \end{matrix}\right\} = \bar{y} \pm t_{f;1-\alpha/2} s_p \left( \frac{(x - \bar{\bar{x}})^2}{s_{xx}} + \frac{1}{K} + \frac{1}{n} \right)^{\frac{1}{2}}$$ **SEq 13**

| Variable | Value |
| --- | --- |
| $\beta_0$ | 38.66645943863722 |
| $\beta_1$ | -3.335984211745958 |
| PCR Efficiency | 0.9941673919755294 |
| $\bar{y}$ | 28.925 |
| $\hat{x}_0$ | 831.9851413821814 |
| $s_p^2$ | 0.00981666666666674 |
| $s_{xx}$ | 14.787116116160096 |
| $\bar{\bar{x}}$ | 3.4889132261647355 |
| $g$ | 0.00031721429593360836 |
| $x_u$ (95% PI) | 757.1503874590653 - 913.763660990488 |
| $\bar{y}_u$ (95% PI) | 28.78882848628346 - 29.061171513716534 |
